# Experimental Directory Structure (Exdir): An alternative to HDF5 without introducing a new file format

**DOI:** 10.1101/249979

**Authors:** Svenn-Arne Dragly, Milad Hobbi Mobarhan, Mikkel Lepperød, Simen Tennøe, Marianne Fyhn, Torkel Hafting, Anders Malthe-Sørenssen

## Abstract

Natural sciences generate an increasing amount of data in a wide range of formats developed by different research groups and commercial companies. At the same time there is a growing desire to share data along with publications in order to enable reproducible research. Open formats have publicly available specifications which facilitate data sharing and reproducible research. Hierarchical Data Format 5 (HDF5) is a popular open format widely used in neuroscience, often as a foundation for other, more specialized formats. However, drawbacks related to HDF5’s complex specification have initiated a discussion for an improved replacement. We propose a novel alternative, the Experimental Directory Structure (Exdir), an open standard for data storage in experimental pipelines which amends drawbacks associated with HDF5 while retaining its advantages. HDF5 stores data and metadata in a hierarchy within a complex binary file which, among other things, is not human-readable, not optimal for version control systems, and lacks support for storing raw data. Exdir, one the other hand, uses file system directories to represent the hierarchy, with metadata stored in human-readable YAML files, datasets stored in binary NumPy files, and raw data stored directly in subdirectories. Furthermore, storing data in multiple files makes it easier to track for version control systems. Exdir is not a file format in itself, but a standard for organizing files in a directory structure. Exdir uses the same abstractions as HDF5 and is compatible with the HDF5 Abstract Data Model. Several research groups are already using data stored in a directory hierarchy as an alternative to HDF5, but no common standard exists in the field. This complicates and limits the opportunity for data sharing and development of common tools for reading, writing, and analyzing data. Exdir facilitates improved data storage, data sharing, reproducible research, and novel insight from interdisciplinary collaboration. With the publication of Exdir, we invite the scientific community to join the development to create an open standard that will serve as many needs as possible and that will serve as a foundation for open access to and exchange of data.

**SIGNIFICANCE STATEMENT:** Experimental Directory Structure (Exdir) is a proposal to standardize a storage solution that has become an increasingly popular alternative to Hierarchical Data Format 5 (HDF5), namely to use directories to define a hierarchy, store data in binary files, and metadata in text files. While this strategy is deployed locally by several research groups, no common standard exists. We envision the establishment of such a standard and present Exdir to the community as a starting point.

## 1 INTRODUCTION

Technology development is continuously driving science to new discoveries. In neuroscience, advancements in genetic tools, recording technology, and computer power have paved the avenue to reveal the underlyings of the healthy and diseased brain. Modern neuroscience usually involves recordings and perturbation at many levels, generating a range of data including imaging, electrophysiology, behaviors, perturbations, and molecular biology. Publication of raw data is acknowledged as critical to enable reproducible research and global large-scale collaborative projects and metadata analyses (Nelson, 2009). However, data from different acquisition systems come in a multitude of data formats that need to be readable for all relevant analysis software and stored for long-term archival. Acquisition systems often use proprietary and specialized formats tailored to data produced by specific types of equipment or software. However, these specialized formats have little applicability outside their intended purpose, making them inaccessible for extended use. In contrast, general-purpose formats can store data for multiple types of equipment and software. When based on open standards, general-purpose formats facilitate data sharing.

Hierarchical Data Format 5 (HDF5)^1^ is a popular and open general-purpose format capable of storing many large and annotated datasets in a hierarchical structure within a single file. HDF5 is the basis of many formats in neuroscience, including the recent collaborative format, Neurodata Without Borders (NWB) (Teeters et al., 2015). However, issues with HDF5 have recently surfaced in the neuroscience community (Rossant, 2016b). Many of these are due to the complex standard of HDF5 and its use of a single binary file to store all the data. The metadata is not human-readable and the binary format is not optimal for version control systems. Further, the use of a single file increases the severity of data corruption, because corruption in a single dataset can affect the entire file. In addition to this, there is no support for storing raw data in other file formats within the HDF5 hierarchy. These issues have sparked a discussion in the wider scientific community on whether HDF5 should be replaced by alternative data formats or if its large feature set outweighs the disadvantages (Hinsen, 2016).

Here, we propose a novel standard, Experimental Directory Structure (Exdir) as an alternative that circumvents the drawbacks of HDF5 and takes advantage of existing, open data formats. Exdir follows the abstract data model used in HDF5^2^, but stores data and metadata in directories to avoid the vulnerability and rigidity associated with storing all data in a single file. Datasets are stored in binary NumPy^3^ files, while attributes and metadata are stored in YAML^4^ text files. Raw data, such as images and time series obtained during data acquisition, can also be stored in Exdir alongside the binary NumPy files. This allows raw data to be organized inside an Exdir hierarchy without any prior conversion, even when the data is composed of multiple formats.

Exdir is ready to use with a reference implementation in Python, a command-line client, and a graphical browser. The application programming interface (API) of the reference implementation is compatible with h5py^5^, a popular HDF5 library for Python. The code is open source and hosted on GitHub^6^.

The idea of an HDF5-replacement based on a hierarchy of directories is already present in the scientific community (Rossant, 2016a), but to the best of our knowledge no formal standard has been introduced. The lack of a standard reduces the opportunities for collaboration through data sharing, and inhibits development of analysis tools. Exdir represents the introduction of a standard that enables novel insight from interdisciplinary collaboration by facilitating reproducible research through improved data storage and sharing. With the publication of Exdir, we invite the scientific community to join the development to create an open standard that will serve as many needs as possible.

## 2 EXISTING ALTERNATIVES

### 2.1 Hierarchical Data Format (HDF5)

HDF5 holds many advantages over alternative data formats, but as described by Greenfield et al. (2015), the HDF5 format also has crucial disadvantages. In the list below, we have summarized the limitations and challenges from Greenfield et al. (2015) that are most relevant to neuroscience along with some additional drawbacks which are addressed with Exdir:

1. Metadata is stored in a binary format which makes it unreadable without tools that read HDF5 files. This also makes the metadata unavailable for text-based command line tools.
2. The specification for HDF5 files is large and complex and there is only one defacto implementation of HDF5 in C that most HDF5-libraries use. Because of the complex specification, this implementation is hard to improve by external contributors. Furthermore, the dependency on one large implementation makes it hard to write software which reads and writes HDF5 files in ways that have not been anticipated by the implementation developers.
3. Like all data formats, HDF5 files are susceptible to data corruption. However, because HDF5 stores all data and metadata in a single file, data corruption in one part of an HDF5 file has a chance of corrupting the entire file.
4. Attributes in HDF5 do not support deeply nested structures, like JSON data, YAML data or Python dictionaries.
5. External version control systems such as Git^7^ and incremental backup systems do not work optimally with HDF5 files because all datasets and metadata are stored in a single binary file. This makes it appear as if the entire file has changed when changes are made to a single dataset.
6. Comparing files in binary formats like HDF5 requires specialized tools. However, text-based formats have a wide range of tools that allow line-by-line comparisons, such as *diff*^8^, and *wdiff*^9^, or the graphical tools *meld*^10^ and *kdiff3*.^11^.
7. Deleting datasets in HDF5 files only removes a reference to the data, while the data remains on disk.
8. Raw data from acquisition or analysis must be stored outside the HDF5 file which makes the raw data detached from the internal hierarchy and inconvenient to annotate. It is often necessary to organize raw data in a separate hierarchy outside the HDF5 file.

### 2.2 Advanced Scientific Data Format (ASDF)

Greenfield et al. (2015) propose a new format (Advanced Scientific Data Format, ASDF) to address many of the above mentioned problems. Similar to Exdir, ASDF also embraces YAML for metadata, but it also stores and organizes binary data in the same YAML file. Storing the data in one file has the same increased risk of data corruption as HDF5 and makes it harder for version control systems to keep track of incremental changes. Furthermore, like HDF5, ASDF does not provide a convenient way to store raw data in the internal hierarchy.

### 2.3 Commonly used open formats in neuroscience

In Table 1, some of the commonly used open formats in neuroscience are listed. Some of these formats are discussed by Teeters et al. (2015) where they also introduce Neurodata Without Borders (NWB), a format recently developed in an attempt to unify cellular-based neurophysiology data and break down barriers for data sharing. Many of these formats, including NWB, are based on HDF5 and therefore share the same advantages and disadvantages as HDF5. Because Exdir is compatible with the abstract data model of HDF5, these formats could be moved to being based on Exdir in the future. The formats that are not based on HDF5 are mostly specialized to neuroscience and therefore have limited applicability to the wider scientific community.

**Table 1.** Overview of commonly used open formats in neuroscience.

| Name | HDF5 | Notes | Reference |
| --- | --- | --- | --- |
| NWB | Yes |  | Teeters et al. (2015) |
| Kwik | Yes |  | Kadir et al. (2014);<br>Rossant et al. (2015) |
| BRAINformat | Yes |  | Rübel et al. (2015) |
| Open Ephys | Yes/No | Binary format specifically designed for electrophysiological data. HDF5 optional | Siegle et al. (2015) |
| NeuroShare | No | Binary format specifically designed for electrophysiological data. | neuroshare.org<sup>12</sup> |
| Neo | N/A | In-memory data format for Python. Uses different formats for file storage. | Garcia et al. (2014) |
| CARMEN NDF | No | Specifically designed for neuroscience. Stores hierarchical structure and metadata in XML files. | carmen.org.uk<sup>13</sup> |
| Nix | Yes | Adds a layer on top of the abstract data model that standardizes annotation of data. Directory-based backend in development. | Stoewer et al. (2014) |
| odML | No | Only applies to metadata. | Grewe et al. (2011) |

### 2.4 Requirements of a new data format

We share many of the requirements reviewed in detail by Greenfield et al. (2015) for the ASDF format. To meet the challenges, a data format should:

1. Have an intrinsic hierarchical structure.
2. Be human-readable.
3. Be based on existing standards.
4. Be easy to extend.
5. Have efficient mechanisms to update data.
6. Have support for both text and binary data. In addition to the above mentioned requirements, we want Exdir to:
7. Minimize the risks and consequences of data corruption.
8. Have a simple, yet flexible specification.
9. Be flexible to data modifications.
10. Be easy to use in ways that have not been anticipated by the authors.
11. Be based on the same abstractions as HDF5 to make it easy to port HDF5-based solutions.
12. Provide a convenient way to store raw data in the same hierarchy.

None of the existing formats known to the authors fulfill all of the mentioned requirements.

## 3 STANDARDS USED IN EXDIR

To fulfill the requirements stated in Section 2.4, we propose a new standard, Exdir, which is based on a standardized directory structure and established open-source file formats. The structure follows the abstract data model used in HDF5, but Exdir uses regular file system directories to define the object hierarchy, and stores datasets, attributes, and corresponding metadata in separate files.

We use YAML to store metadata and attributes. YAML is a human-readable format and has the ability to store many types of objects, provided software to convert the object to and from YAML. Furthermore, libraries for YAML support exist for most major programming languages, including Python, C/C++, Java, and Rust^14^.

To store binary data we have chosen the NumPy format. This is a simple, efficient, and widely used file format. Furthermore, there exist libraries to load NumPy files in several languages such as Matlab^15^, Rust^16^, R^17^, and C/C++^18^.

The hierarchical structure of Exdir benefits from the hierarchy of directories in file systems. It is an existing standard which is familiar to computer users. By using this inherent hierarchy, Exdir makes it possible for a user to browse any Exdir object with a native file explorer. Further, the use of regular directories allows raw data from acquisition to be stored in the same hierarchy and annotated together with the rest of the data.

Parallel reading and writing to different objects in an Exdir directory is handled by the operating system. This is in contrast to HDF5, where parallel read/write operations must be handled by the HDF5 library because all datasets are stored in the same file. Parallel read/write operations to the same dataset is currently not supported in the reference implementation of Exdir, but is planned for a future release or extension.

As each dataset is stored in its own directory, the risk of data corruption is reduced. If one dataset is corrupted, it is unlikely to affect the other files in a directory. This separation also makes Exdir avoid the problem of data remaining after deletion in HDF5 and taking up space. Deleting a dataset in Exdir immediately frees up disk space.

When accessing large Exdir `File` objects, one can easily retrieve and share subtrees of the main hierarchy by copying the corresponding directories. This reduces memory, CPU, and server-communication costs by keeping the size of data handled to a minimum. When sharing Exdir data with others, one can use readily available compression file formats such as `.zip` or `.tar.gz`.

Instead of building data consistency verification into Exdir, we envision the use of existing tools for this purpose. For instance, plugins (see section 5.2) can be developed to make version control systems like Git track the version of each object in an Exdir directory and ensure that no files have changed independently. This also allows Exdir to be combined with Git-based systems like Gin^19^, which are tailored towards cloud-based handling of large datasets (Garbers et al., 2017).

## 4 BASIC STRUCTURE OF EXDIR DIRECTORIES

Exdir has four types of objects, `File`, `Group`, `Dataset`, and `Raw`, where each is represented as a directory in the file system. `Raw` is a type of object that is not present in the original HDF5 abstract data model. Metadata for each of these objects is stored in a file named `exdir.yaml`.

All objects can have attributes stored in an optional file named `attributes.yaml`. Figure 1 shows an example structure of an Exdir `File`, and a summary of specifications of the data format is shown in Table 2.

**Figure 1.**
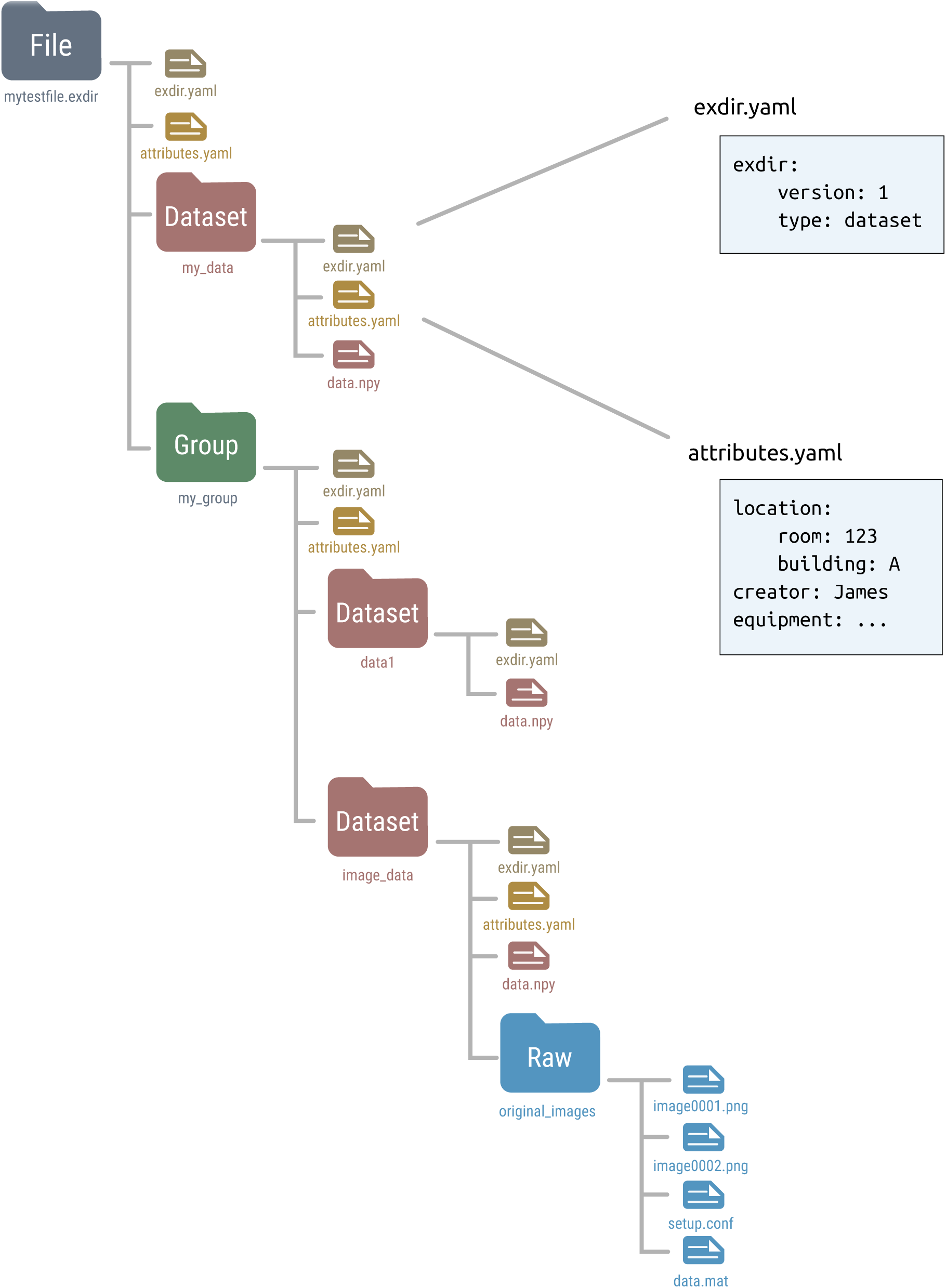
Overview of an example Exdir directory. `File, Group`, and `Dataset` refer to objects in Exdir, and are stored as directories in the file system. These objects are equivalent to the same objects in the HDF5 abstract data model. `Raw` is specific to Exdir and is a regular directory containing arbitrary data files. Inside each directory, there is a file named `exdir.yaml` with information about the object type and Exdir version. Each object may contain an `attributes.yaml` file containing user-defined attributes. Inside the `Dataset` directory is a file named `data.npy` that contains the data of the dataset stored in the NumPy binary format.

**Table 2.** Exdir format structure.

| Type | Description | Contains | Required | Optional |
| --- | --- | --- | --- | --- |
| File | Root object | Group<br>Raw<br>Dataset | `exdir.yaml` | `attributes.yaml` |
| Group | Intermediate directory | Group<br>Raw<br>Dataset | `exdir.yaml` | `attributes.yaml` |
| Dataset | Data |  | `exdir.yaml`<br>`data.npy` | `attributes.yaml` |
| Raw | Arbitrary data files |  |  | `exdir.yaml`<br>`attributes.yaml` |

### 4.1 Metadata, attributes, and data files

Metadata for each object is stored in the `exdir.yaml` file in the object’s directory. This file defines that the current directory is an Exdir object, and contains information about the Exdir version and object type. For example, this is the `exdir.yaml` file of a dataset:

~~~
exdir :
    version : 1
    type: dataset
~~~

The object type can either be `file`, `group`, `dataset`, or `raw`. The `exdir.yaml` file is optional for `Raw` objects.

User-defined attributes of an Exdir object are stored in that object’s directory in the `attributes.yaml` file. Attributes are stored as key–value pairs, which can be nested:

~~~
location :
    room: 123
    building : A
creator : James
equipment : …
~~~

Binary data of a `Dataset` is stored in the NumPy format^20^ in a file named `data.npy` in the `Dataset` object’s directory.

### 4.2 File

The `File` object is the root (top level) object of an Exdir hierarchy. Every directory below a `File` in the directory hierarchy is part of that `File`. A `File` cannot contain other `File` objects. The metadata of the `File` is stored in `exdir.yaml`, and optional attributes in `attributes.yaml`.

### 4.3 Group

Inside the `File`, multiple objects may be stored, among them `Group` objects. `Group` objects may also contain any number of other `Group` objects, `Raw` objects, and `Dataset` objects. `Group` objects are stored as directories in the file system with metadata stored in `exdir.yaml`, and optional attributes in `attributes.yaml`. File objects are a specialization of a `Group` object.

### 4.4 Dataset

`Dataset` objects are for storing data. `Dataset` objects are stored as directories with metadata in the `exdir.yaml` file, and user-defined attributes in an optional `attributes.yaml` file. The data within a `Dataset` is stored in a binary NumPy file named `data.npy`.

### 4.5 Raw

`Raw` objects are used to store data in other formats than the NumPy format. While the user may store any type of data in the a `Raw` directory it is encouraged to use `Dataset` objects if possible. For `Raw` directories the `exdir.yaml` file is optional. Further, attributes are stored in the optional `attributes.yaml` file. There is no similar concept of `Raw` objects in HDF5.

## 5 REFERENCE IMPLEMENTATION IN PYTHON

We have created a reference implementation of the Exdir standard in Python. This implementation is hosted on Github and is publicly available with an open-source license. It can easily be installed with Anaconda^21^.

The reference implementation of Exdir owes its relative simplicity to being based on existing formats, and to having a hierarchy based on regular file system directories. It is implemented using the open-source NumPy and PyYAML^22^ libraries, and is designed to be compatible with the popular HDF5 library, h5py. The compatibility should ease the transition from h5py to Exdir.

The class hierarchy of the reference implementation is shown in Figure 2. The `Raw, Group`, and `Dataset` classes inherit from `Object`, which contains their common methods. The `File` class is a subclass of `Group` and they share many of the same methods. `Attribute` is a separate class that handles attributes for all Exdir objects. Furthermore, the reference implementation has an extensive test suite that can be run with `pytest`^23^.

**Figure 2.**
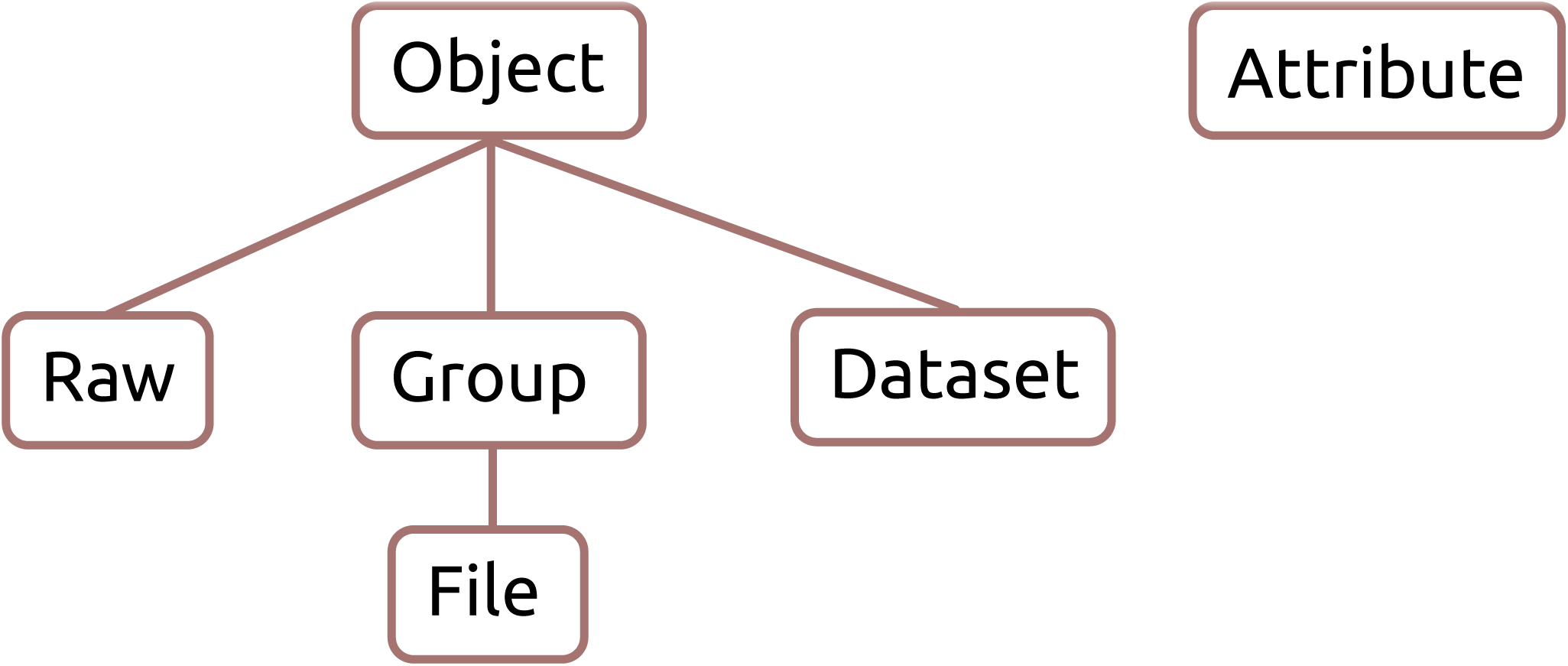
Exdir reference implementation class hierarchy.

### 5.1 Overview of the Exdir API in Python

In this section we give a quick overview of the Exdir Python API. An Exdir `File` is created as follows:

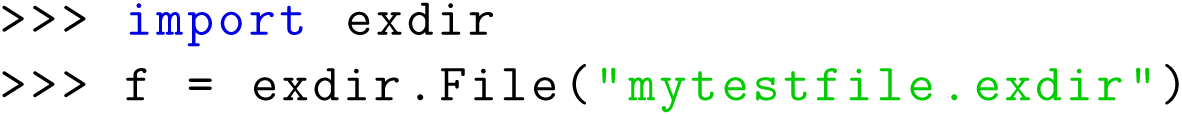

The `File` object points to the root directory in the Exdir directory structure. To create a `Dataset` inside the root directory (or other `Group` objects) the `create_dataset()` method can be used:

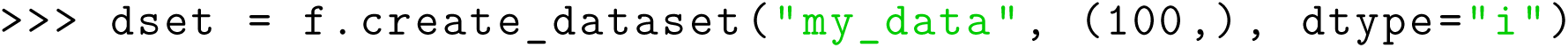

Exdir `Dataset` objects are not NumPy arrays, but behave similarly. They have both a shape and a data type:

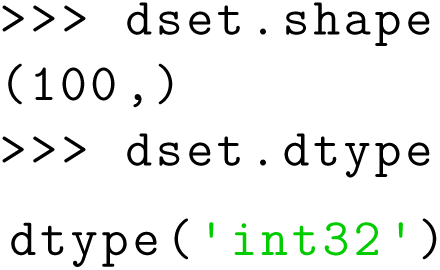

`Dataset` objects support array-style slicing, which can be used to read and write data to the `Dataset`:

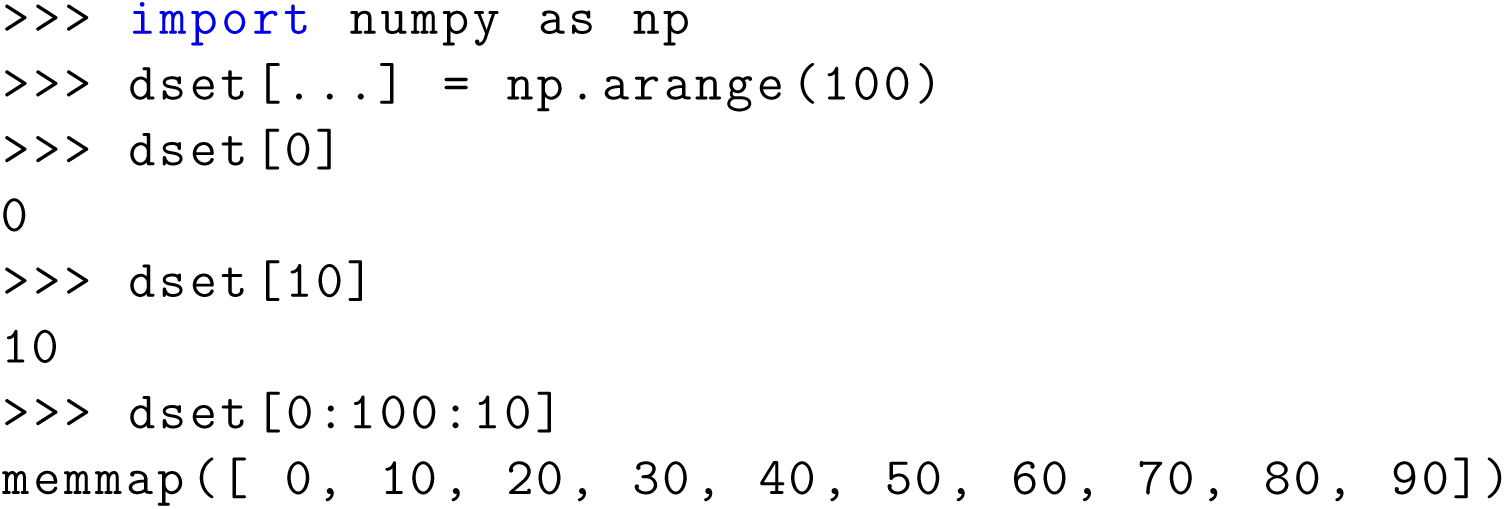

In addition, `Dataset` objects can also be created from the data directly:

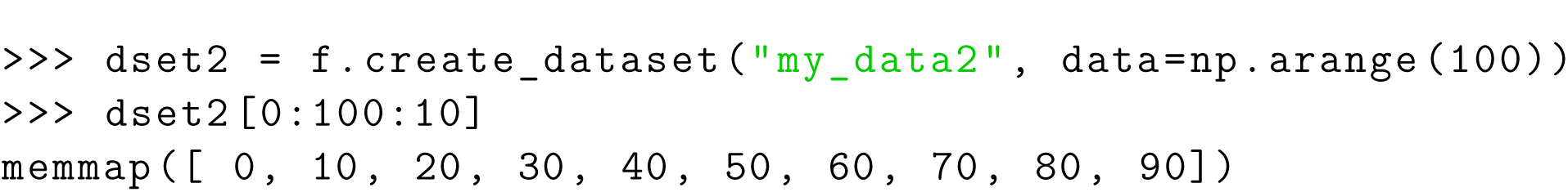

Exdir uses NumPy’s memory mapping feature (memmap) to access segments of larger datasets on disk, without reading the entire file into memory. Furthermore, Exdir supports all the operations supported by memmap, including fancy indexing:

~~~
>>> dset[dset [:] > 90]
array([91 , 92, 93, 94, 95, 96, 97, 98, 99] , dtype=int32)
~~~

An Exdir `Group` can be created using `create_group()`:

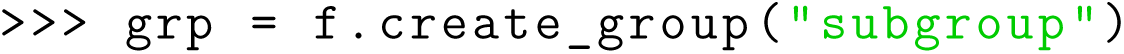

As with `File` objects, a `Dataset` is created inside a `Group` by using the `create_dataset()` method:

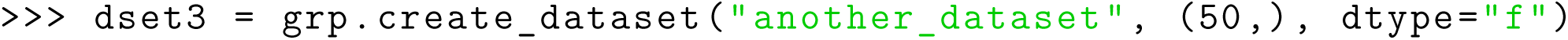

`Group` objects support most of the Python dictionary-style interface. You retrieve objects in the file using the item-retrieval syntax:

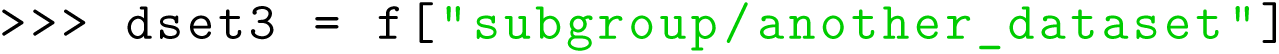

As shown above the name of objects follows the hierarchy of the POSIX standard with /-separators. To retrieve the name of any object in an Exdir directory one can use:

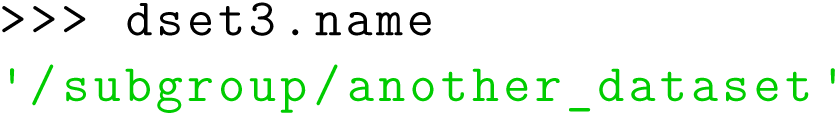

Iterating over a `File` or a `Group` provides the names of their members:

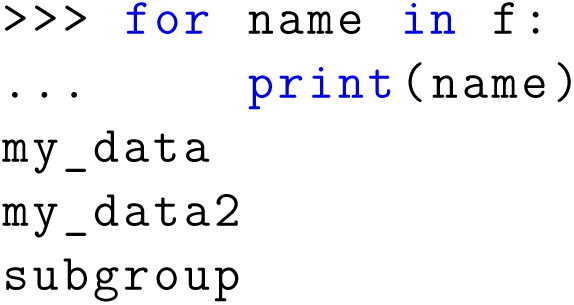

Containership testing also uses names:

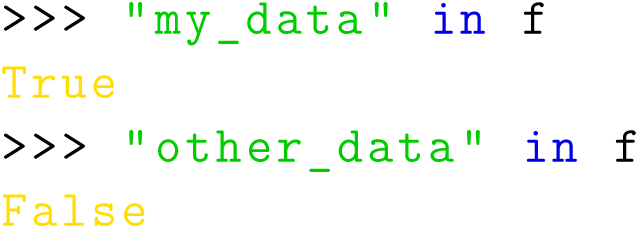

`Group` objects have the methods: `keys(), values(), items(), iter()`, and `get()`.

All `File` objects, `Group` objects, and `Dataset` objects can have attributes. Attributes are accessed through the `attrs` property, which implements a dictionary interface:

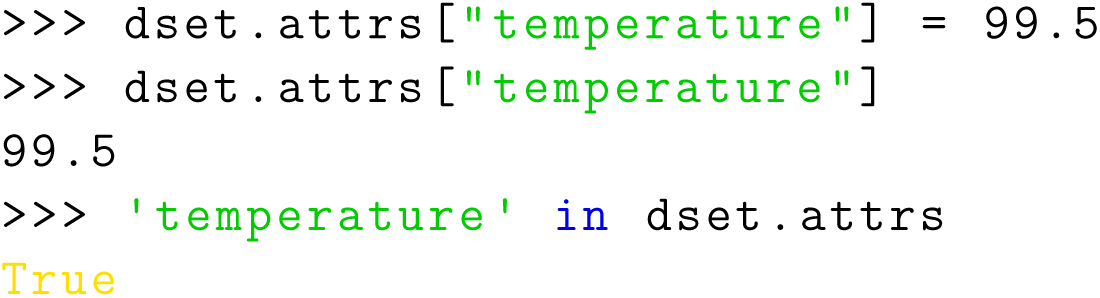

Unlike HDF5 and h5py, Exdir supports dictionaries as attributes:

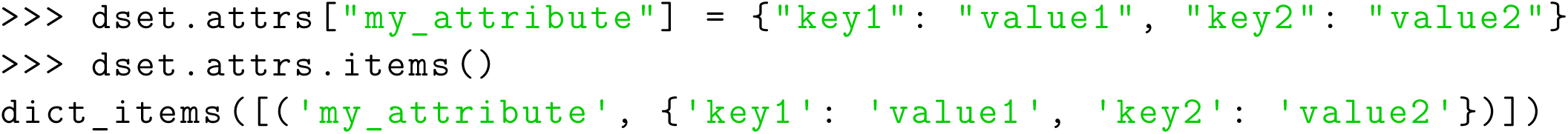

After the above commands, the Exdir directory structure becomes:

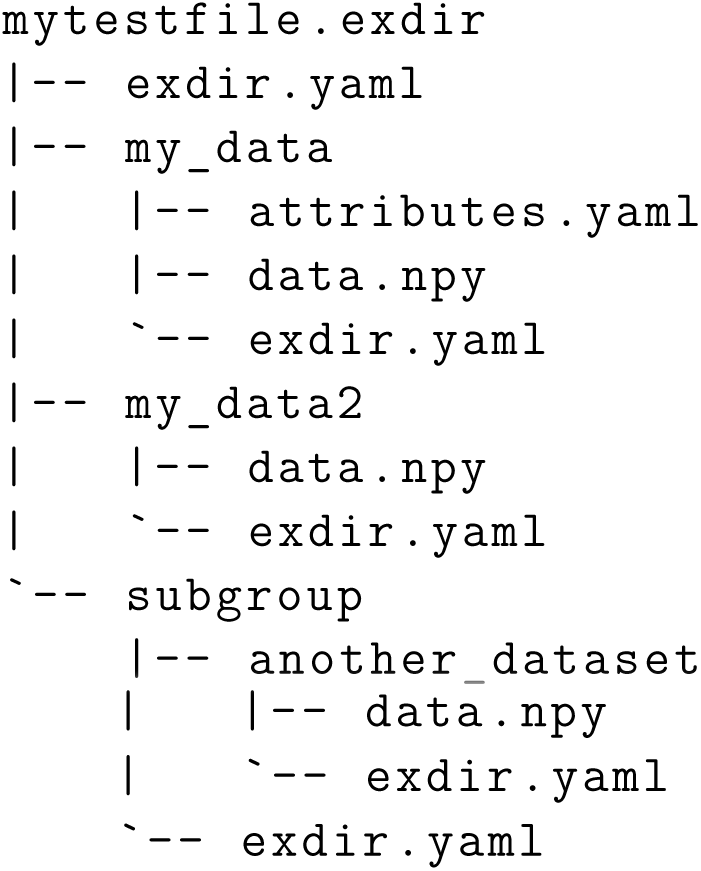

### 5.2 Exdir Plugins

The functionality of Exdir can be extended with plugins. These allow modifying the behavior of Exdir when enabled. For instance, dataset and attribute plugins can perform pre- and post-processing of data during reading and writing operations. Some plugins are provided in the `exdir.plugins` module, while new plugins can be defined by Exdir users or package developers.

One of the built-in plugins provides experimental support for units using the `quantities` package^24^:

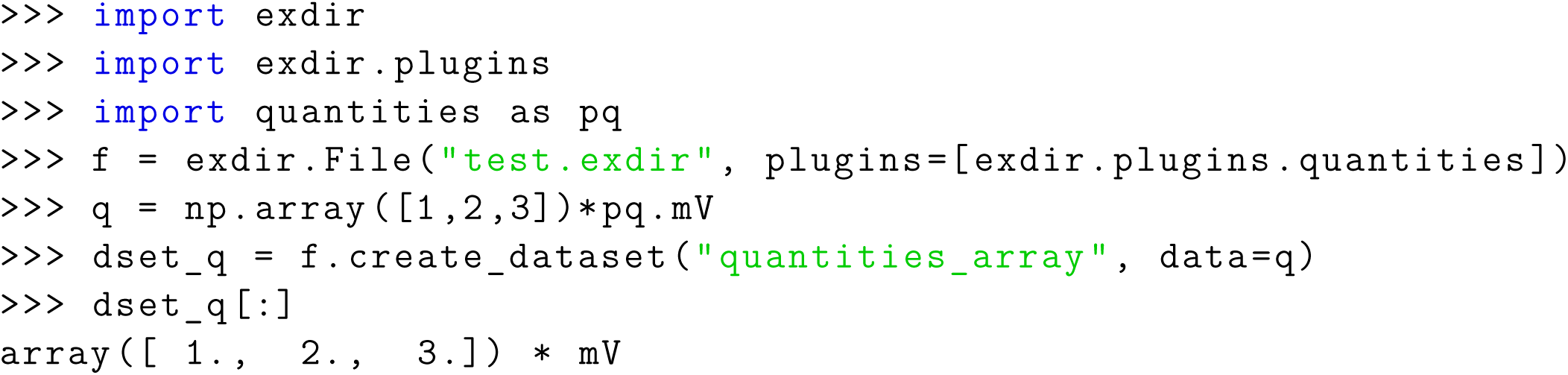

As shown in the above example, a plugin is enabled when creating a `File` object by passing the `plugin` to the plugin argument.

To create a custom plugin, one of the handler classes in `exdir.plugin_interface` must be inherited. The abstract handler classes are named after the object type you want to create a handler for. The following is an example of a `Dataset` handler that multiplies the numbers in the `Dataset` by 2 whenever the `Dataset` is written to file:

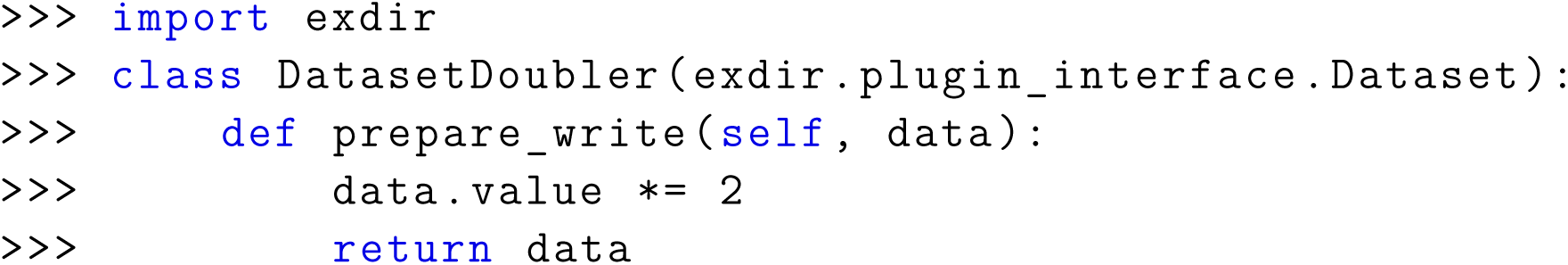

In the example above, `data` is an object that has properties `value, attrs`, and `plugin_meta`. The property `attrs` is a dictionary with optional attributes, while `plugin_meta` is a dictionary with information about the plugin.

We create a plugin that uses this handler as follows:

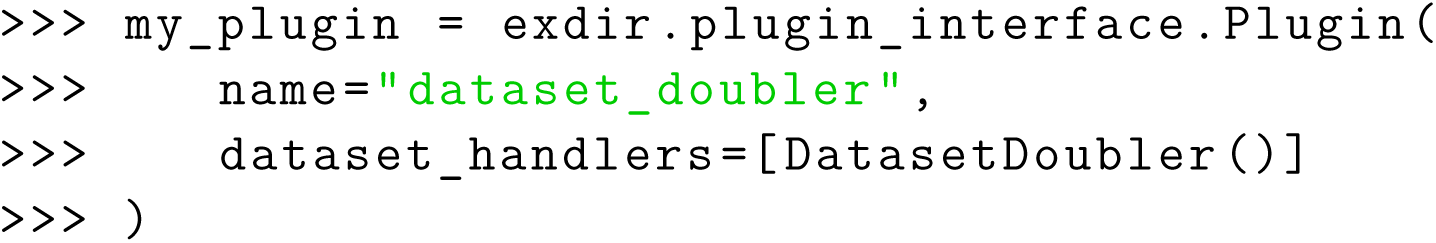

The plugin is enabled when opening a `File` by passing it to the `plugins` parameter:

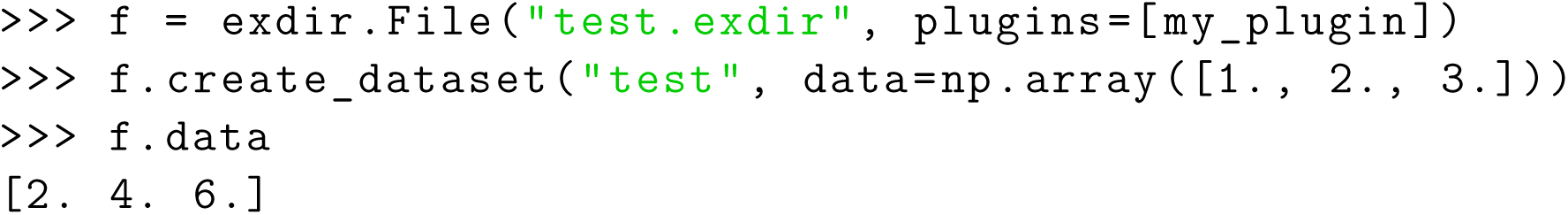

### 5.3 Converting from hdf5 to Exdir

As can be seen from Table 1, many common formats in neuroscience are based on HDF5. Since Exdir follows the abstract data model of HDF5, it is easy to switch from HDF5 to Exdir, and these formats should be able to support both HDF5 and exdir as backends. Often, the only change needed to transition from h5py to Exdir will be to switch from the following:

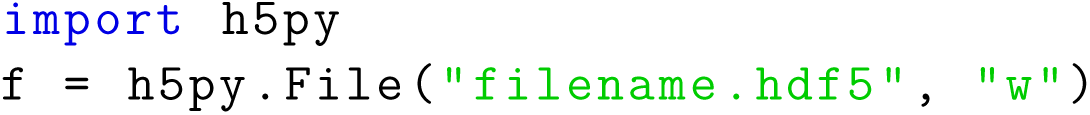

To the following:

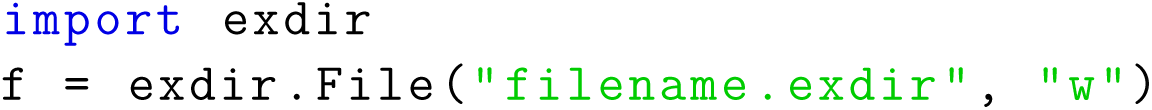

In most cases, the rest of the code can be left unchanged.

A few operators in h5py are missing in the reference implementation and will eventually be added. Further, HDF5 has support for linking of objects, which is currently not part of the Exdir specification and will be added in the future. Finally, the reference implementation currently does not support parallel read/write operations on single files. A future plugin is planned to provide such support.

## 6 TOOLS FOR EXDIR

The Exdir command line interface and the Exdir browser are tools created to make it easier to work with Exdir data.

### 6.1 Exdir command line interface

Exdir-cli is a simple command line interface for browsing Exdir directories and to create Exdir `File` objects and `Group` objects. Listing the content of an Exdir `File` is done in the command line by the following:

~~~
$ exdir list mytestfile . exdir
group1
group2
dataset
~~~

Listing the contents of a Dataset is done by the following:

~~~
$ exdir show dataset
nums
Type: Dataset
Name: /dataset
Shape : (23632 ,)
Data:
[0 0 0 …, 6 6 6]
~~~

### 6.2 Exdir browser

Exdir browser is a graphical user interface for viewing and editing Exdir directories (see Figure 3). The browser can be installed on Linux, macOS, and Windows through Anaconda^25^ or from source^26^.

**Figure 3.**
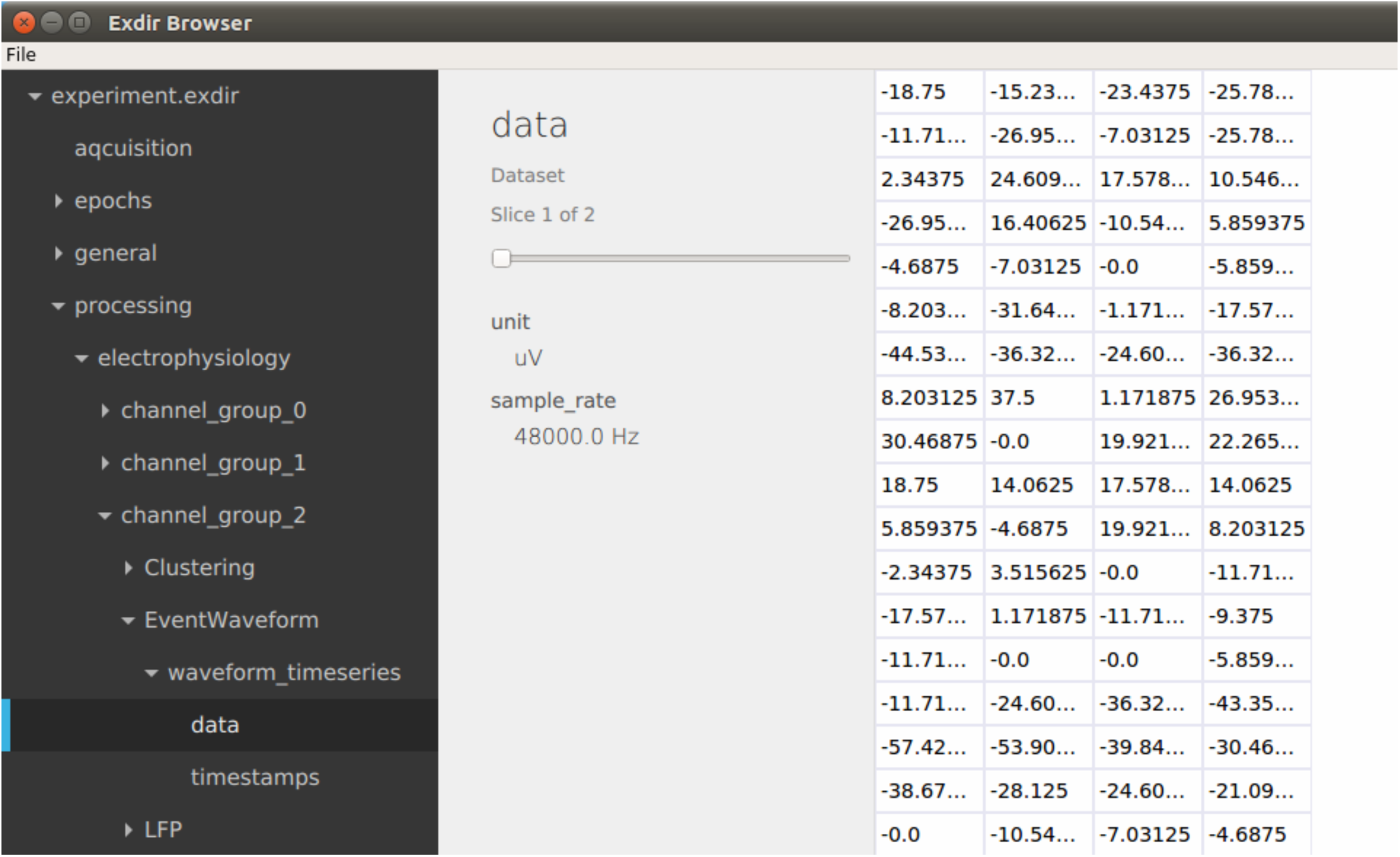
Screenshot of the Exdir browser.

After opening an Exdir directory, the Exdir browser shows a hierarchical tree of all the objects in that directory. Information about each object is shown when selected and attributes of all objects may be edited. `Group` objects can be expanded to show their child objects, similar to directories on the file system. When selecting a dataset, the contents is shown in a 2D table. If the dataset is three dimensional, you can select the slice.

## 7 PERFORMANCE

As with other formats, the performance of Exdir is limited by the file system and underlying hardware. In general, data readability has been prioritized over performance in Exdir, but we are improving the performance where possible.

We have performed benchmarks for some common operations and compared the Exdir reference implementation to the h5py Python library. The results are listed in Table 3. These benchmarks were performed on an in-memory virtual hard disk (RAM disk) to obtain more reliable results. Note that this only gives an indication of the performance differences between h5py and the Exdir reference implementation, as not all aspects of running on a physical hard drive are reflected.

**Table 3.** Results from benchmarks comparing performance in Exdir with h5py. A 2GB RAM disk was used as virtual hard drive for the tests. Software used: Python 3.6, NumPy 1.13.1, Ubuntu 16.04. Hardware used: Intel Core i7-4820K 3.70GHz, 32 GB RAM.

| Test | Exdir | h5py |
| --- | --- | --- |
| Add 5 attributes | 0.0044 s | 0.00086 s |
| Add 100 attributes | 0.11 s | 0.0048 s |
| Add dataset with $10^6$ 64-bit floats | 0.0088 s | 0.0040 s |
| Add dataset with $10^8$ 64-bit floats | 0.16 s | 0.17 s |
| Create 5000 groups | 52 s | 0.24 s |
| Create tree (3 groups × 5 levels) | 2.73 s | 0.079 s |
| Read/write 3D slice ($100 \times 100 \times 100$) | 0.0048 s | 0.0033 s |

As can be seen from table 3, metadata manipulation in Exdir is slow compared to HDF5. This is because we have chosen to store metadata in human-readable YAML text files. This is a deliberate choice we made because we believe readability of these files is more important than performance. However, manipulation of metadata in Exdir can outperform HDF5 on networked file systems if the file system downloads and uploads entire files when they are modified. Metadata in Exdir is stored in separate files, and only these files need to be downloaded, while the rest of the dataset can remain on the server. This is in contrast to HDF5 where the entire file may have to be downloaded.

Reading and writing large continuous data in Exdir is about as fast as in HDF5. This is also the case for reading and writing to parts of a dataset. However, HDF5 supports storing chunked data, which is a feature missing in Exdir, and in these cases, HDF5 is likely to outperform Exdir when reading and writing binary data.

Creating many empty objects performs poorly in Exdir, while it is fast in HDF5, as shown in the Create 5000 groups benchmark in Table 3. This is because Exdir creates a directory and an `exdir.yaml` file for each object, while HDF5 only needs to add a reference to the new object in the main file.

In summary, the performance of Exdir is mostly limited by the performance of the file system and the performance of the YAML and NumPy libraries. Exdir performs worse than HDF5 with many small objects, but performs similarly when working with larger datasets. Furthermore, Exdir performs worse than HDF5 with many read/write operations on metadata. We therefore recommend using a dedicated database if metadata operations are frequent and become a performance bottleneck.

## 8 DISCUSSION

We have proposed a new standard, Exdir, that puts the abstractions of HDF5 on top of a hierarchical directory structure. Exdir gives the same flexibility as HDF5, but with the advantages of a simpler specification, human-readable metadata, and applicability of established tools. Further, the hierarchy and metadata can be modified manually without tools specific to Exdir, while the data is accessible by existing libraries for common languages. This makes Exdir a possible replacement for HDF5 in computational and experimental data pipelines.

We have presented a reference implementation in Python, a command-line client, and a graphical browser that are all open source and available on GitHub. Together, these tools will hopefully make it easy for other researchers to explore the standard and provide valuable feedback. Because Exdir is based on the established NumPy and YAML formats, we expect APIs for other languages to be fairly easy to implement.

The reference implementation has an extensive test suite and has been thoroughly tested, although the format is still under development. The flexibility of the format gives many possibilities for future development. Exdir includes the concept of plugins, which makes it easy to extend implementations with new functionality without adding more complexity to the specification.

Because similar strategies for data storage are already in use, but no formal standard exists, we believe Exdir provides an opportunity for increased data sharing and development of tools that can be shared across multiple disciplines. We hope Exdir can lay the foundation for a standardization of such strategies, and contribute to the general discussion on data storage in science.

## AUTHOR CONTRIBUTIONS

SD, MHM, and ML conceived of and designed the project. SD, MHM, ML, and ST wrote software, documentation, and the paper. All authors contributed to revising the paper and approved of the final version.

## ACKNOWLEDGMENTS

The development of Exdir owes a great deal to other standardization efforts in science in general and neuroscience in particular, among them the contributors to HDF5, NumPy, YAML, PyYAML, SciPy, Klusta Kwik, NeuralEnsemble, and Neuroscience Without Borders.

## Footnotes

1 https://hdfgroup.org

2 https://support.hdfgroup.org/HDF5/doc/

3 http://numpy.org

4 https://yaml.org

5 http://h5py.alfven.org

6 https://github.com/CINPLA/exdir/

7 http://git-scm.org

8 https://www.gnu.org/software/diffutils/manual/diffutils.html

9 https://www.gnu.org/software/wdiff/manual/wdiff.html

10 http://meldmerge.org/

11 http://kdiff3.sourceforge.net/doc/index.html

14 http://yaml.org/

15 https://github.com/kwikteam/npy-matlab

16 https://github.com/potocpav/npy-rs

17 https://github.com/eddelbuettel/rcppcnpy

18 https://github.com/rogersce/cnpy

19 https://web.gin.g-node.org/

20 https://docs.scipy.org/doc/numpy/neps/npy-format.html

21 https://anaconda.org/cinpla/exdir

22 http://pyyaml.org

23 https://docs.pytest.org

24 https://github.com/python-quantities/python-quantities

25 https://anaconda.org/cinpla/exdir-browser

26 https://github.com/CINPLA/exdir-browser

